# Maternal antibodies and density dependence affect suppression of host populations by novel pathogens

**DOI:** 10.1101/2025.09.12.675835

**Authors:** James J Bull, Rustom Antia

**Affiliations:** Dept of Biological Sciences, University of Idaho, Moscow, ID USA; Dept of Biology, Emory University, Atlanta, GA USA

## Abstract

Current theory for the regulation of host populations by pathogens suggests that a high level of suppression during the initial epidemic phase will be followed by a population rebound with decreased virulence due to pathogen and host evolution, and the extent of host suppression increases with increasing pathogen transmissibility (*R*_0_) and virulence. Using simple epidemiological models, we explore the effect of two factors on short- and long-term suppression: the strength of density-dependent population regulation (homeostasis) and maternal antibodies. We showed previously that, in the absence of maternal antibodies, the strength of homeostasis can greatly effect long term population suppression. Here we find that maternal antibodies can significantly reduce suppression of the host population if they attenuate rather than block infections, but then only for rapid homeostasis. A higher *R*_0_ can result in lower suppression, and the average virulence can decline over time without any (genetic) evolution. Our results suggest the need for a nuanced view of long-term suppression by a new pathogen, with the outcome sensitive to many details even in the absence of evolution.

## Introduction

Viruses and other pathogens can cause massive mortality when first introduced into naive animal or human populations [1, 21]. The introduction may be a deliberate attempt to suppress the host population as with myxomavirus and caliciviruses to control Australian rabbits [8, 11]. Or a pathogen may naturally jump from its reservoir host into a new host, as with the 1918 human influenza virus in humans and distemper viruses in lions and in seals [7, 28, 30, 31]. Finally, a pathogen can be introduced via contact with previously isolated populations of the same species, as happened to the Aztecs following contact with Spanish conquistadors [9, 22].

Considerable attention has been devoted to the initial epidemic or pandemic spread of the pathogen and its effect on the host population, as illustrated in the examples mentioned above. Other studies have considered the long-term impact of a pathogen on its host population. These ecologically motivated studies considered the role of pathogens in regulating host populations. In a classic study, Anderson and May [1] showed that, since the transmission (*R*_0_) of the pathogen is proportional to the density of hosts, suppression of the host population is greatest when the density of the host is high.

Many studies that considered how the long term impact of pathogens differ from the intial impact focus on evolution. Evolutionary changes, whether in the pathogen or host, are difficult to predict, but the general view (supported by the myxoma virus released in Australian rabbits) is that an initially highly lethal pathogen will evolve to be less virulent and the host population will evolve resistance.

There are, however, non-evolutionary mechanisms that can change the long term impact of the pathogen from its initial impact, and these have received relatively less attention than pathogen evolution. One factor is a reduction in the average age of infection as the pathogen becomes endemic; the virulence of infections is often lower for youth than for adults [18, 20, 23]. Long term suppression may also be considerably reduced if young are protected by maternal antibodies [12, 32]. Both of these factors can reduce the severity of infections as the disease transitions to endemicity, and for pathogens used to control animal populations, can allow the host population to rebound.

We use mathematical models to explore the impact of a pathogen in suppressing the host population and how this changes between the early, epidemic stage and the late, endemic stage. We focus on non-evolutionary mechanisms and specifically consider the consequences of viral transmissibility (*R*_0_) and maternal antibodies. We show that the extent of density-dependent regulation in the host population, which we term homeostasis, has an unexpectedly large effect on the ability of maternal antibodies to suppress the host population in the long-term. Our findings are relevant to the use of pathogens to control pests as well as to conservation efforts to mitigate natural epidemics. A major goal is to keep the models simple enough to allow the analysis of model structure and the effect of specific parameters on outcomes. As such, the study is not an attempt to match any particular application.

## Results: Maternal antibodies do not invariably enable population recovery

Maternal antibodies, transferred from the mother to offspring, are known to protect the young for a few months after birth [5, 10, 32]. Maternal antibodies have been shown to protect young rabbits from myxoma virus infection [10, 19], and some consequences of this protection to population control have been modeled [12, 13, 14]. Maternal antibodies might provide protection in two ways: by **blocking** – reducing the susceptibility of individuals to becoming infected – or by **attenuation** – reducing the severity of infections while enabling long-lived immunity [16, 19, 27].

Intuition suggests that if maternal antibodies attenuate the infection, considerable population recovery might ensue from an otherwise highly lethal pathogen. This recovery would start from the few naive individuals who survive an infection who then have offspring with maternal antibodies. If those offspring are infected during the period of protection, the attenuation improves their survival, resulting in high levels of antibodies with which they endow on their progeny. If levels of infection are high, this benefit can be propagated as if inherited epigenetically – a type of Lamarckian inheritance. A protected class of the population may thus emerge and expand disproportionately because their survival is much higher than of naive individuals.

The consequences of maternal antibodies that block infection are not so clear: offspring are temporarily blocked from infection but may be infected later in life – suffering the typical case mortality (by assumption here). If blocking is lifelong, the recovered individual is protected for life, but its progeny will not be protected. So the effect of blocking by maternal antibodies may be to reduce the number of susceptible individuals in the population and thereby lower the incidence of the pathogen. But blocking is not inherited because a blocked individual cannot be infected and thus does not develop any phenotype to be passed on.

We use a simple compartmental model that incorporates susceptible (*S*), infected (*I*) and recovered (*R*) populations. The model allows for individuals protected by maternally transferred antibodies by either of two mechanisms: progeny with maternal antibodies may be protected from infection or experience attenuated infections of reduced severity. We therefore model the effect of maternal antibodies by introducing two additional classes of individuals: *S*_*M*_ are individuals born with maternal antibodies and thus protected, and *I*_*M*_ are individuals who became infected while in the *S*_*M*_ category. The equations for maternal antibodies are thus:

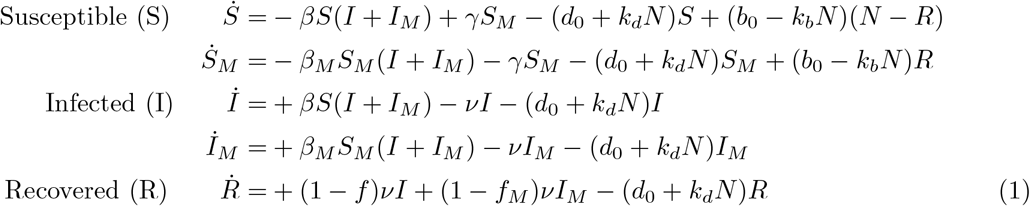

where notation is given in Table 1.

**Table 1:** Notation for variables and parameters in eqns (1). Two different sets of values for the four birth-death parameters are used, one represeting rapid homeostasis, the other slow homeostasis. Birth and death parameter values are chosen so that the equilibrium population size in the absence of the pathogen is 1, and the parameters for rapid homeostasis are chosen to represent a short-lived, high-fecundity species such as rabbits (see Supplementary file S1 for details); values for slow homeostasis are chosen to contrast with those for rapid homeostasis.

| Symbol | Description | Value |
| --- | --- | --- |
| $S$ | Susceptible, born of naive mothers | densities, $\leq 1$<br>functions of time |
| $S_M$ | Susceptible with maternal antibodies | |
| $I$ | Infected from $S$ | |
| $I_M$ | Infected from $S_M$ | |
| $R$ | Recovered and immune | |
| $N$ | Total population | |
| $\nu$ | loss of infection rate | 50 / year |
| $\beta, \beta_M$ | transmission rate constants | for $R_0$ ($1.5 < R_0 < 10$) |
| $f, f_M$ | case mortality | $0 < f < 1$ per infection |
| $\gamma$ | maternal antibody decay rate | 4 / year |
| Rapid homeostasis |  |  |
| $b_0, d_0$ | intrinsic birth and death rates | 3.5, 0.2 /year |
| $k_b, k_d$ | parameters for logistic growth | 2.5, 0.8 |
| Slow homeostasis |  |  |
| $b_0, d_0$ | intrinsic birth and death rates | 1.5, 0.8 /year |
| $k_b, k_d$ | parameters for logistic growth | 0.5, 0.2 |

In the absence of the pathogen, the population size is maintained by density dependence of the birth and death rates, logistic growth. We let the birth rate decrease linearly with density [*b*(*N* ) = *b*_0_ − *k*_*b*_*N* ] and the death rate increase linearly with density [*d*(*N*) = *d*_0_ + *k*_*d*_*N*]. Through appropriate choice of parameter values, the carrying capacity (equilibrium population density) in the absence of pathogen is here always scaled to 1. Density dependence also operates when the pathogen is present, the consequence being that births exceed deaths except for disease-caused mortality.

The severity of infections is described by the case mortality (*f* ). We choose case mortality rather than a virulence rate [e.g., 15], as it gives a more intuitive description of virulence than a rate of virulence, and it allows exploring the effect of independently changing transmission and disease severity without affecting the lifespan of infecteds. The basic reproductive number (*R*_0_) equals *β/*(*d*_0_ + *k*_*d*_ + *ν*) ≈ *β*/*ν*, an approximation that relies on the duration of infection (1/*ν*) being much shorter than the lifespan of the host (1/(*d* + *k*_*d*_)). The simulations that vary *R*_0_ do so by changing transmission, *β*. *β* and *f* are subscripted with *M* to accrue to hosts protected by maternal antibodies.

We use parameter values to represent a small mammal, such as a rabbit, which might apply to the well documented example of Australian rabbits and other invasive mammals. Two scenario’s are addressed for host population regulation in the absence of the pathogen. In the first scenario, the population exhibits rapid regrowth when culled, which we refer to as *rapid homeostasis*. Rapid homeostasis is modeled by choosing high values for *k*_*b*_ and *k*_*d*_, the parameters that describe the density dependence of birth and death rates. The effect of *slow homeostasis* is modeled by assigning smaller *k*_*b*_ and *k*_*d*_ values so that the birth and death rates exhibit a smaller dependence on population size. The pathogen-free equilibrium is unaffected, however.

How do maternal antibodies affect population suppression, and how robust are the effects? We consider separately that maternal antibodies block infections or that they attenuate infections – reduce lethality but allow the generation of long-term immunity. Blocking is modeled by setting *β*_*M*_ = 0, attenuation by setting *f*_*M*_ = 0 and *β*_*M*_ = *β* for the protected individuals. These are extremes and are chosen to maximize the possible effects. The parameter for waning of protection from maternal antibody is set as *γ* = 4, which corresponds to protection for an average of 3 months. Each analysis of maternal antibodies is run in parallel for rapid and for slow homeostasis, as previous analyses found that homeostasis had a large effect in the absence of material antibodies [3].

### Short-term suppression is unaffected by maternal antibodies

The initial level of population suppression – the suppression experienced at the end of the epidemic phase – is largely independent of homeostasis and of maternal antibodies. This insensitivity to homeostasis arises because the epidemic occurs on a much shorter time scale than do births and deaths. The initial level of suppression in the absence of maternal antibodies has been studied extensively as a theoretical problem – by assuming that births and deaths can be ignored – and the magnitude of the short-term epidemic is known as the epidemic ‘final size’ [2, 17, 24, 25]. In our model, the extent of population suppression in this initial phase is simply the final size times case mortality. Likewise, maternal antibodies have minimal effect on initial suppression because the initial suppression occurs prior to the appearance of individuals with maternal antibodies. This insensitivity to homeostasis and maternal antibodies can be inferred from Fig. 1, where it is seen that the initial drop is essentially the same for rapid and slow homeostasis and regardless of maternal antibodies (panels A, B, for specific parameter values). Panel (C) offers a heatmap based on the final size calculations, where it is easily seen that the initial suppression is strongly affected by case mortality (*f*) and that suppression increases strongly with *R*_0_ when *R*_0_ is small but becomes largely insensitive to *R*_0_ at higher values. It is also clear from panels (A) and (B) that this approximate equivalence between rapid and slow homeostasis and independence of maternal antibodies is short-lived.

**Figure 1.**
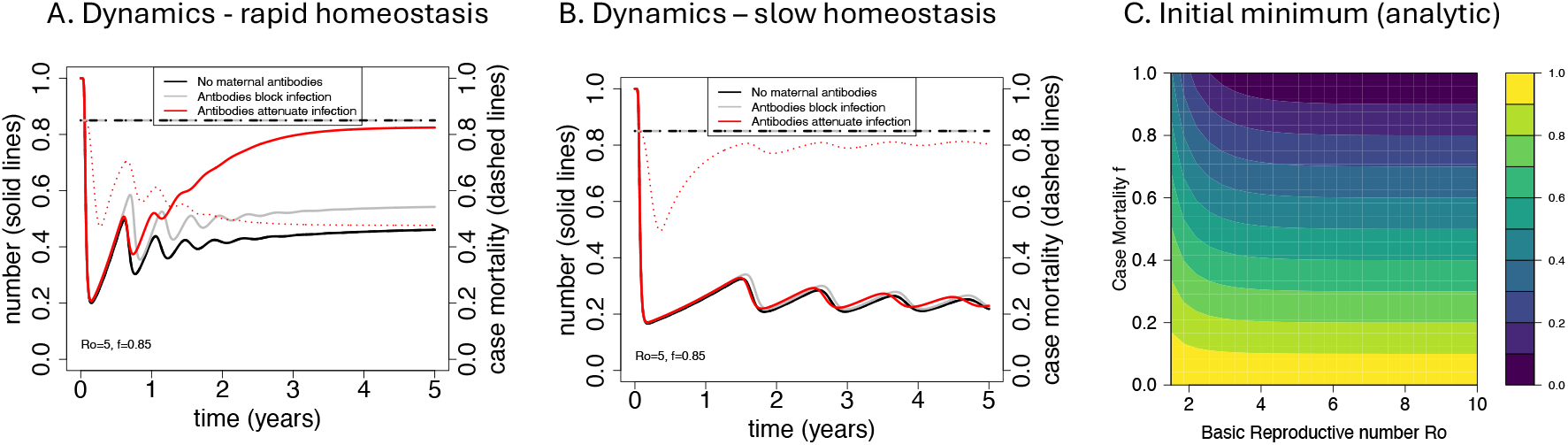
Dynamics of populations under (A) rapid and (B) slow homeostasis, and (C) a heatmap of the initial population minima per different pathogen *R*_0_ and case mortality (*f* ). Assuming specific parameter values, panels (A) and (B) each contrast the dynamics for no maternal antibodies (black), maternal antibodies block the infection (gray) and maternal antibodies attenuate the infection (red). The initial minima are unaffected by homeostasis or maternal antibodies, but beyond the initial minima, the population rebounds substantially only for rapid homeostasis when maternal antibodies attenuate. The heatmap (C) shows that the initial minima are strongly affected by case mortality (*f* ) and also somewhat by *R*_0_. Dashed curves in (A) and (B) give average case mortalities, which change when maternal antibodies attenuate the infection, because there is then a class of infections that are protected from dying. Parameter values are given in Tables 1, and *R*_0_ = 5 and *f* = 0.85 for panels (A) and (B). The heatmap in (C) is based on analytical calculations of the epidemic ‘final size’ given in the text.

The short-term dynamics beyond the initial epidemic phase in panels (A) and (B) foreshadow results below, for steady states: maternal antibodies attenuating infections results in a striking recovery of the host population, yet this happens only when homeostasis is rapid (red curve in panel A). When homeostasis is slow or maternal antibodies block infection, there is little to no additional rebound compared to an absence of maternal antibodies.

### Long-term recovery is possible from maternal antibodies but depends on biology

The long-term outcome will differ from the short-term one at least because birth and death rates influence the long-term population sizes. But there will also be time for maternal antibodies to develop and be transmitted to young. However, it is not guaranteed that maternal antibodies will enable significant recovery. First, if the effect of maternal antibodies is to block the infection, population recovery may be minor (given the reasoning offered above). Second, maternal antibodies will wane in progeny, so if the parasite is not common in the population, most infections may occur after maternal antibodies have waned and no longer protect. While it is easy to appreciate the qualitative effects of these processes, it is challenging to anticipate the quantitative effects without formal analysis.

Fig 2 presents panels illustrating several aspects of our analysis, always contrasting rapid and slow homeostasis (top and bottom rows, respectively). All panels are heatmaps for different combinations of *R*_0_ and case mortalities (*f*) at steady state. In order, the columns are: (i) population sizes in the absence of maternal antibodies for comparison to steady-state population sizes when maternal antibodies (ii) block or (iii) attenuate infection, respectively. (iv) Population-wide, average case mortality at endemicity for the case of maternal antibody attenuation. The case mortality *f* on the vertical axis applies to naive infections, whereas the population average includes attenuated infections protected by maternal antibodies, which here are assumed to have a case mortality of 0.

**Figure 2.**
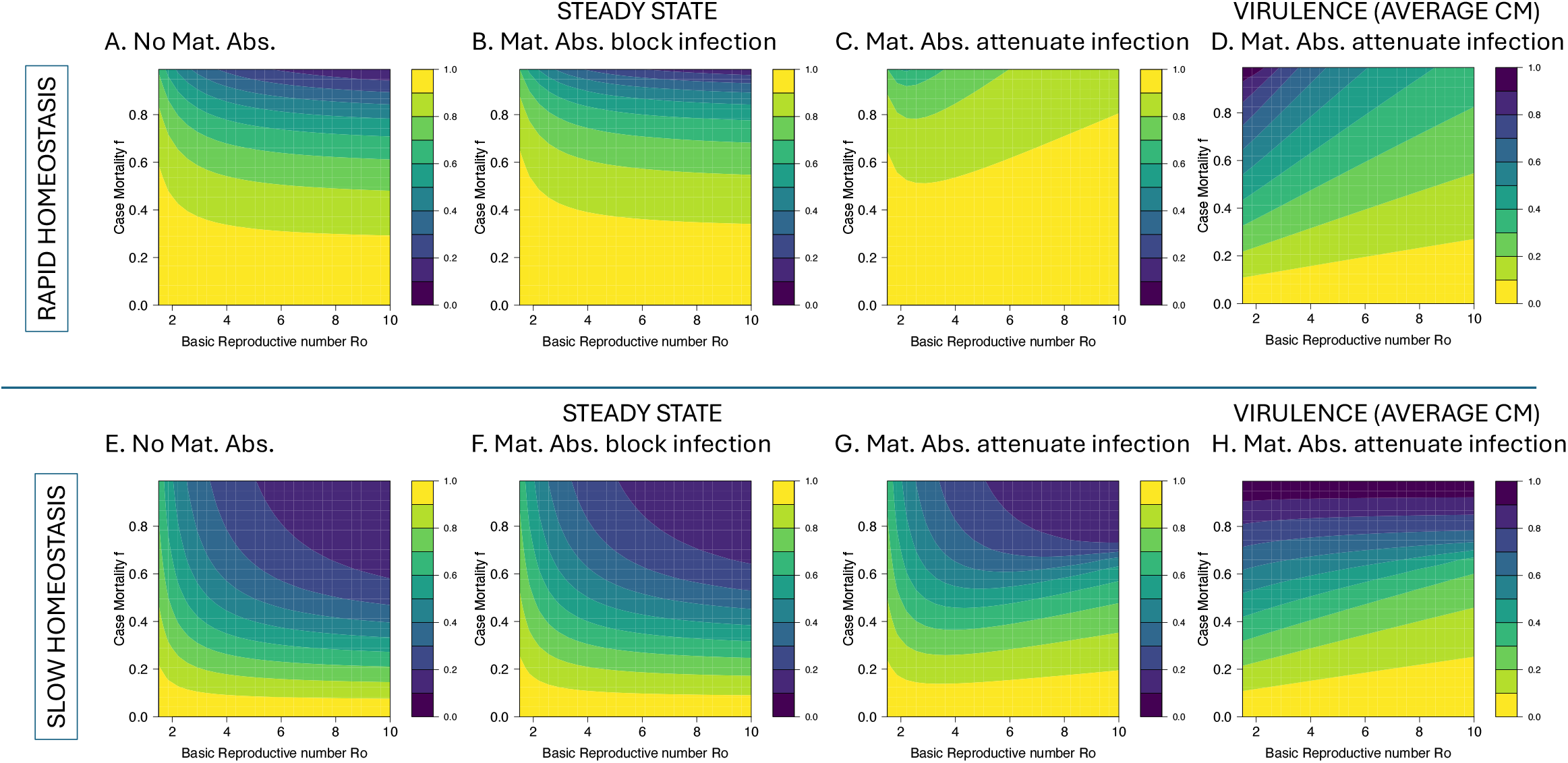
Heatmaps showing the effect of maternal antibodies on host population suppression and virulence. The top row is for rapid, the bottom row for slow homeostasis. All panels are for steady states. Panels (A) and (E) in the first column provide long term population sizes in the absence of maternal antibodies. Population suppression is greater for slow than for rapid homeostasis (A-C versus E-G), also greater for blocking than for attenuating the infection (B versus C, and F versus G), although this difference is much greater for rapid than for slow homeostasis. The rightmost panels (D, H) show the reduction in average case mortality at steady state when maternal antibodies attenuate infection; the upward-sloping contours indicate that average case mortality declines with *R*_0_. Parameter values are given in Tables 1; *R*_0_ = 5 and *f* = 0.85 for panels A and D. The rate of waning of maternal antibody protection *γ* = 4 which corresponds to protection for 1/4 year.

There are several strong patterns in the Figure. (i) There is a large effect of homeostasis on all long term outcomes. Rapid homeostasis promotes far greater population recovery than does slow homeostasis – the populations in panels (A)-(C) are larger than those in (E)-(F). The explanation thus transcends any effect of maternal antibodies: in suppressed populations, net births are higher under rapid homeostasis than slow homeostasis [3]. (ii) There is little effect of blocking (compare panels A to B and E to F). (iii) Maternal antibodies attenuating infections results in a striking, long term recovery of the host population, but this happens only when homeostasis is rapid.

Any benefit of maternal antibody attenuation can be realized only when offspring are infected prior to losing maternal antibodies. Thus we expect – and observe in panels (C) and (G) – the paradoxical effect that population recovery benefits from high *R*_0_ because the higher transmission leads to a higher incidence of infection and thus earlier infection.

### Antibody duration

A higher *R*_0_ benefiting population recovery with maternal antibody attenuation was explained on the grounds that higher *R*_0_ results in infections occurring at earlier ages – prior to the waning of protection by maternal antibodies. Similar reasoning suggests that any effect of maternal antibodies on host populations will depend on antibody duration – 1*/γ* in our model. If antibodies wane too fast relative to the average age of infection, then few or no individuals will be protected when they are eventually infected. In turn, average age of infection will depend on *R*_0_ and on the birth and death rates at steady-state. Thus, the strong population recovery observed for rapid homeostasis under attenuation as well as the broad lack of recovery under slow homeostasis may change, depending on antibody biology.

Consequences of changing the duration of protection by maternal antibodies are shown in Fig. 3. As expected, increasing antibody duration increases long term population size under attenuation for both rapid and slow homeostasis (panels B, D). A similar but milder trend is also seen for blocking with rapid homeostasis (A), but there is only a very slight effect of antibody duration evident for blocking with slow homeostasis (C). As before, host population suppression is greater for slow homeostasis than for rapid homeostasis.

**Figure 3.**
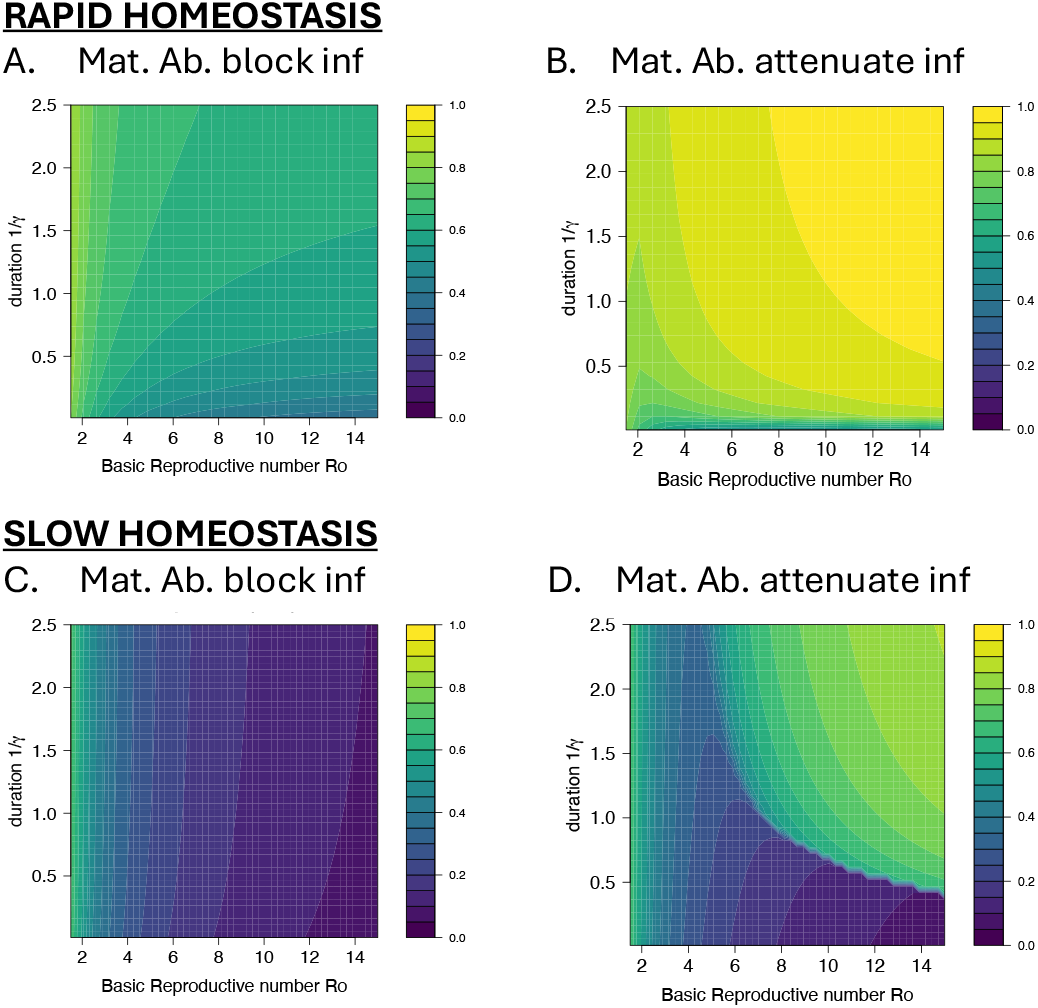
Effect of maternal antibody duration on the suppression of the host population at steady state. Plots vary *R*_0_ (X-axis) and the duration of protection by maternal antibody (1*/γ*, Y-axis). As above, suppression is typically greater for slow homeostasis than for rapid homeostasis. The effect of antibody duration is moderately weak except in panel D and at very short durations. The top row is for rapid homeostasis, bottom is for slow homeostasis. Case mortality (*f* ) is 0.85, and the intrinsic lifespan 1*/d*_0_ = 5 years.

**Figure 4.**
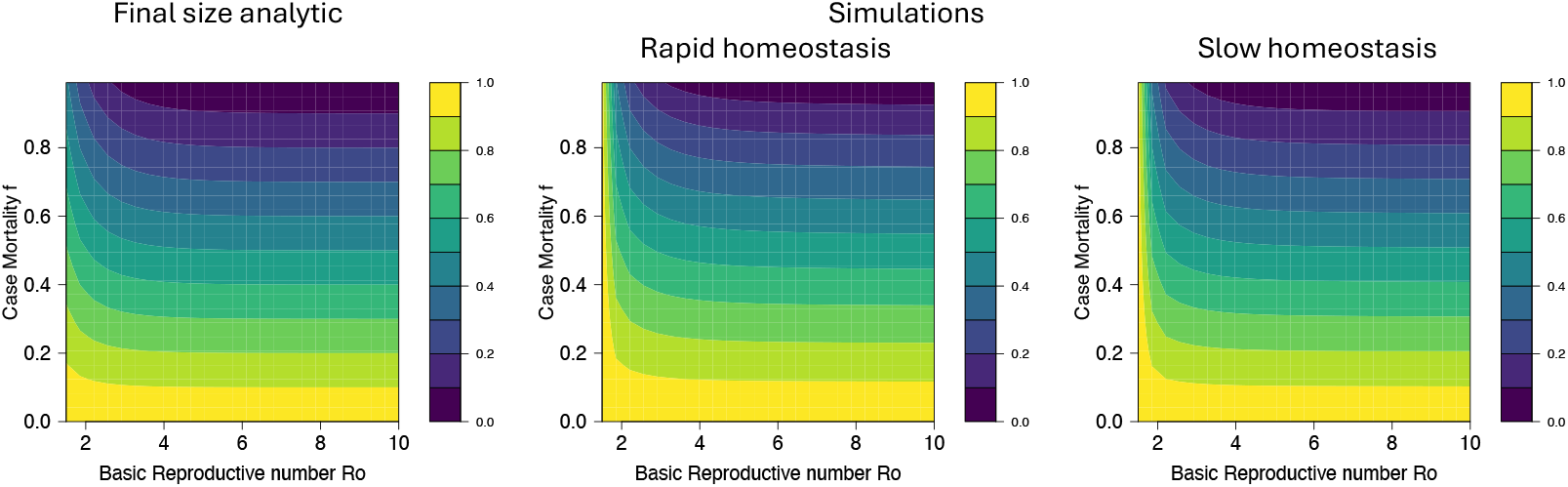
For the basic model (1), plots compare the analytic approximation for the initial minimum (left) and the actual initial minimum (middle and right). They are virtually indistinguishable except at the left margin. The discrepancy at low *R*_0_ arises because, for these *R*_0_, the pathogen spreads slowly and allows significant numbers of births during the initial phase.

**Figure 5.**
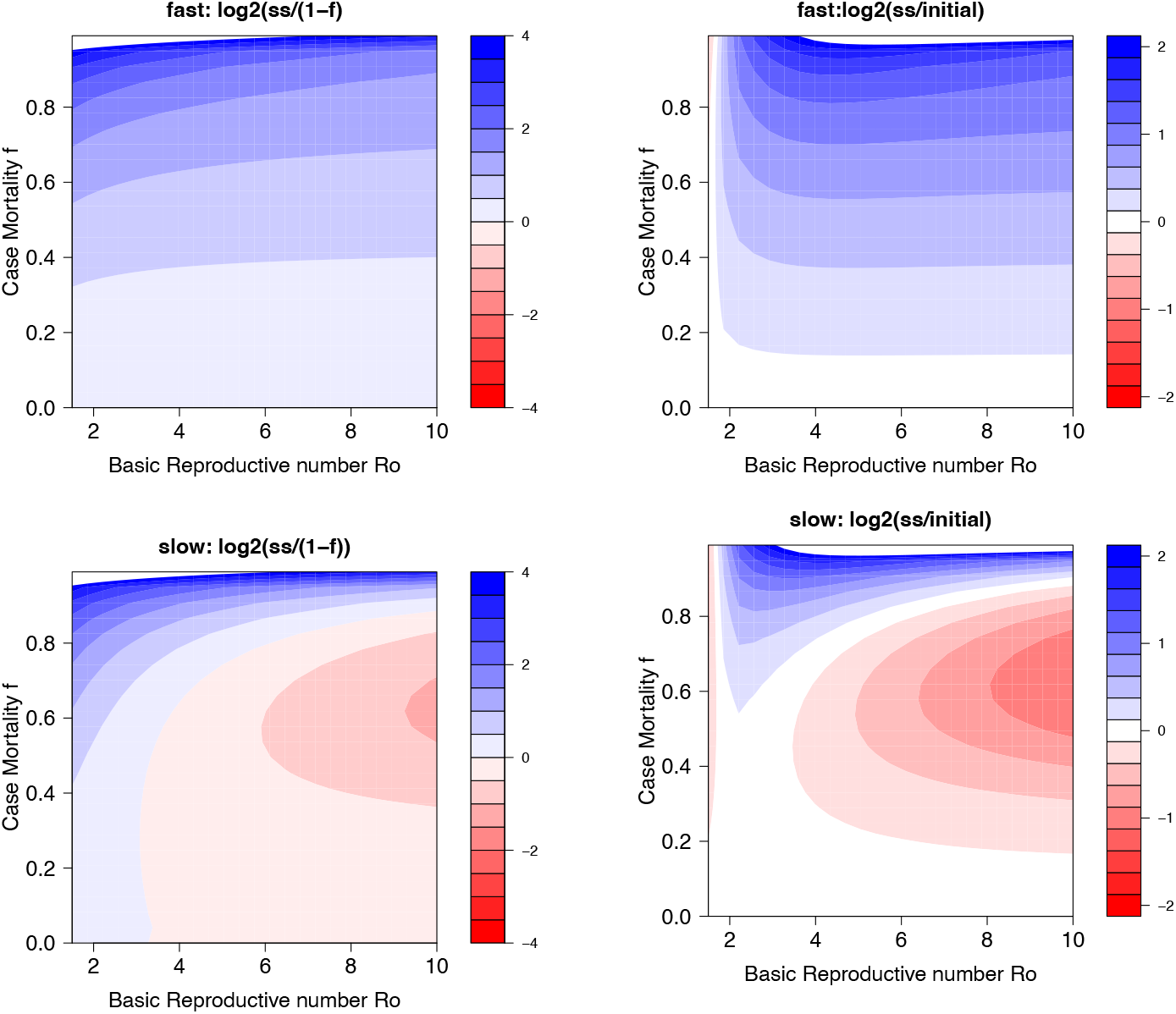
Population suppression at steady state can sometimes exceed case mortality, but not at the initial minimum. Red/pink shows the zones in which population size is less than 1 − *f*, which is the fraction of cases surviving infection. Thus, red and pink show the zones in which the population is suppressed beyond the level that would be imagined from experimental studies of case mortality. Top plots show the log2(Steady-state/initial minima). Bottom plots show log2(steady-state/(1-f)). Parameter values are the same as for text Fig. 1.

Fig. 3 provides some interesting insights to blocking. Reinforcing the impression from Fig. 2 that blocking had small effects on long term outcomes, panels A and C of Fig. 3 also reveal subtle, mild effects of antibody duration with blocking – which implies that maternal antibody blocking is affecting long term population size, if only slightly. Furthermore, the effect of *R*_0_ is in the opposite direction as with attenuation – higher *R*_0_ leads to increased suppression. As case mortality is the same for all infections under blocking, higher *R*_0_ must have this directional effect because it leads to more infections.

Perhaps the most interesting result is the sharp boundary observed in panel D: with slow homeostasis and maternal antibody attenuation: population suppression is minimal until reaching a threshold combination of *R*_0_ and antibody duration, beyond which population size rebounds greatly. Although the overall trend is the same for rapid homeostasis (panel B), it is the sharp boundary in panel D that is of interest. A fine-grained analysis of steady state values reveals an extremely sharp transition in all state variables, one that is sensitive to initial conditions. Whether such thresholds would operate under realistic conditions is debatable, but their presence in the idealized context of the model may portend difficulties with prediction in the field, with slight variations in *R*_0_ or antibody duration shifting the outcome from success to failure of population control.

### Non-genetic reductions in virulence with attenuation

When maternal antibodies attenuate infections, there will be two levels of case mortality, that of naive infections and that of protected infections. Since mortality of infections in individuals with maternal antibodies is low (0 in our simulations), any increase in the incidence of these infections will result in a decrease in the population-wide case mortality – a reduction in the virulence of the pathogen. When the pathogen first invades, the average case mortality per infection (virulence) is *f* – because maternal antibodies are initially absent and all infections are naive. As the prevalence of individuals with maternal antibodies increases there is a reduction in the *average* lethality of infections.

Fig. 2 panels D and H show the average case mortality at steady state. The value on the vertical axis gives the average case mortality when all infections are naive, as when the pathogen first invades. From the upward slopes of the contours, it is easily seen that the average case mortality at steady state decreases with increases in *R*_0_. This decrease is because as *R*_0_ increases there is a decrease in the age of infection and thus a higher probability that individuals with maternal antibodies (*S*_*M*_) get attenuated infections prior to waning of their maternal antibodies. The patterns are different between rapid and slow homeostasis – extreme reductions at high *f* for rapid homeostasis, but for slow homeostasis, virtually no reductions at high *f* despite obvious reductions at low *f* .

The dashed red curves in Fig. 1 panels (A) and (B) show the time course of changes in average case mortality over five years along with changes in population size . There is no obvious association of virulence with changes in population size, and in panel B, there is an early major reduction in virulence despite almost none in the long term.

These virulence reductions are not evolutionary, but they could easily and mistakenly be interpreted as evolutionary because they occur as the pathogen invades and becomes established. The reduction in virulence is greatest when the *R*_0_ of the pathogen is high because in this regime, many individuals with maternal antibodies get mild infections before the protection from antibodies wanes.

If maternal antibodies block infection, there is no change in the average virulence of infections – because blocked individuals do not get infected, and once blocking has waned, infections are fully virulent. The dashed black lines in Fig. 1 panels (A) and (B) show these non-changing virulences merely for comparison to the attenuation case.

## Discussion

Our study used mathematical models to investigate the impact of a pathogen on the population size of its host. The pathogen might be deliberately introduced to control an invasive species, as with rabbits in Australia and France [11, 19], or a pathogen might inadvertently jump to a new species of economic or conservation interest. This study builds on the pioneering work of Kermack and McKendrick as well as of May and Anderson [1, 21] that considered the role of pathogens in regulating host populations. Their papers focused on how pathogen regulation of the host population depends on the nature of the pathogen – its mode of transmission and the duration of host immunity. That work established some principles that have since become the standard understanding: (i) for a given level of disease-induced mortality, the extent of population suppression increases with increases in the pathogen’s basic reproductive number (*R*_0_), and (ii) the population typically rebounds after a high level of initial suppression.

Analyses here explored this question by considering factors that might alter the long term outcome: density-dependent population regulation and protection of offspring by maternal antibodies. In a previous study [3], we considered changes in the rate of density-dependent population regulation with no maternal antibodies. We observed the outcome described by earlier studies that population suppression increases with increases in both *R*_0_ and case mortality. We found that at steady state, population sizes for the same pathogen could differ substantially, depending only on the strength of density-dependent regulation intrinsic to the population.

The current study introduced one biologically-inspired property expected to affect population suppression: maternal antibody protection of progeny. Our analysis suggests that the population-protective effect of maternal antibodies is much greater if they do not prevent infection but attenuate its severity. This allows individuals who get infected while having maternal antibodies to acquire long-term protective immunity and to endow their progeny with the same protection. If maternal antibodies attenuate infections there is a qualitative change in how increases in *R*_0_ affect the population size and average case mortality, but chiefly for rapid homeostasis. Having a higher *R*_0_ could result in an *increase* in the population size at steady-state; this arose because a higher *R*_0_ led to an increase in the fraction of individuals with maternal antibodies getting mild, immunizing infections prior to the loss of these antibodies.

This work follows the precedents of three Fouchet et al. papers [12, 13, 14], who tailored their models to myxoma virus in rabbits and included several details that we intentionally omitted: seasonal reproduction by the rabbit plus maternal antibodies affecting three infection properties at once (blocking, attenuation, transmission). As here, Fouchet et al. [12] found that maternal antibodies were protective of the rabbit population at high transmission rates but not at low (their analysis was limited to a 100-fold difference in *R*_0_). We too found the protective population effect of maternal antibodies to be sensitive to transmission as well as other details, although our analyses evaluated each property separately: blocking infection versus attenuation of case mortality, antibody duration, *R*_0_ and density-dependent homeostasis. These combined results suggest that any prediction of maternal antibody effects in a field context will be difficult. Our results highlight the importance of understanding the mode of action of maternal antibodies – specifically whether they block vs. attenuate infections, and indeed whether there is a transition from blocking to attenuation as the titer of maternal antibodies wanes over time.

When maternal antibodies attenuate, average case mortality (‘virulence’) declined between the time of pathogen invasion and endemism. This change is not evolutionary in a strict sense, rather it reflects changes in the relative abundance of benign versus lethal infections as the pathogen becomes common. Those changes in virulence superficially resemble evolution of reduced virulence. If the pathogen was to die out and then be reintroduced, the process would repeat.

Efforts to control invasive rabbits with viruses in Australia and France should in principle offer relevant case studies for evaluating these models [11, 19]. However, there are important omissions from our simple models that complicate comparisons to rabbit data: ongoing reinfection by myxoma virus so that immunity is not lifelong [19] and seasonal reproduction of rabbits [29]. In France, geographic variation in rabbit suppression from myxoma virus suggests that local factors are important [19]. Those factors could reflect structural omissions from our models or could stem from local variation in parameter values such that our models would match local abundances when parameterized correctly. Regardless, it seems clear that general models such as those here will not be appropriate for specific applications without modification. At the same time, the models point to principles that may apply widely and be important components in any specific application.

Although many aspects of the Australian efforts to suppress rabbits were carefully documented [11], quantitative details of the magnitude of population suppression do not appear to be readily available. Two viruses were introduced: myxoma virus, followed decades later by a calicivirus causing rabbit hemorrhagic fever disease. One survey spanning the introduction of myxoma virus in 1950 to 2013 shows a rapid 90% suppression of the rabbit population with only a small rebound over 3-4 decades [Fig 6 in 6]; a stylized plot from a different study of rabbits in South Australia shows a much faster recovery (until fleas were introduced) from an equally impressive initial suppression [Fig. 1 in 26]. Details of both surveys are minimal. If maternal antibody protection was operating, the rebounds should have been far more rapid than is evident in those surveys. Fouchet et al. [12] commented that young rabbits lacked viral receptors for myxoma virus, which would be consistent with blocked infections in young (not due to maternal antibodies), but a lack of viral receptors is inconsistent with another paper reporting early infection of rabbits [19]. The data from those studies are thus fragmentary and do not provide much basis for a test of the models.

**Figure 6.**
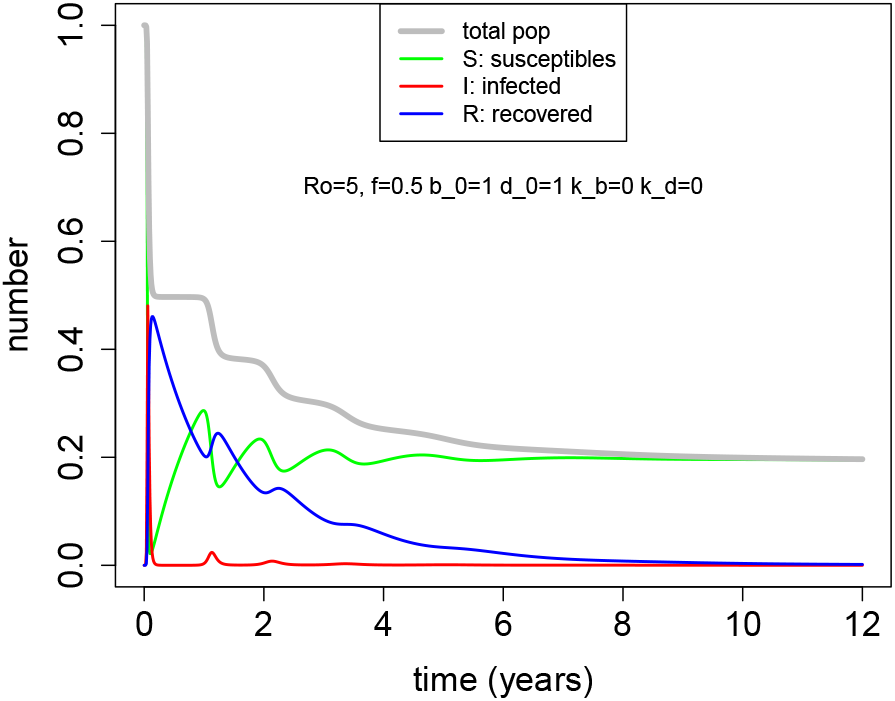
In the absence of homeostasis the total population declines until the reproductive number falls below 1. The early process involves stepwise decreases in population size, each preceded by an increase in the number of susceptibles that eventually enables the pathogen to expand further; recovereds decline as the susceptibles increase.

Our study leaves unexplored many obvious extensions of the models. Some specific extensions are relevant to the field observations mentioned above: seasonal host reproduction and stochastic (non-deterministic) viral abundance, model properties which might accommodate viral extinctions. More generally, what properties of density dependence are important for long term suppression or recovery? Our model allowed 4 birth and death parameters and presented analysis of only two sets of values (although we have documented the same kinds of effects for other sets of values). Analysis of more parameter combinations will shed light on just which properties of density dependence are important. Indeed, our preliminary trials suggest that a model with a single density-dependent parameter may reproduce the qualitative results reported here. Furthermore, it will be desirable to consider different model structures for density dependence than the one we applied (which was linear in population size). Another important consideration is that our model decoupled case mortality and recovery from infection. A more common approach is to assign mortality and recovery rates from infection, whereby a high mortality rate shortens the duration of infection and thus reduces *R*_0_ [4, 15]. If the model structure of case mortality greatly affects suppression, then it adds a further challenge in predicting the long term impact of a lethal pathogen. Last, we are aware that age-dependent differences in case mortality can have large effects on long term suppression.

## Acknowledgments

We acknowledge the following support: JB: P20-GM104420; RA: U01 AI150747 and U01 AI144616.

## File S1: Supplementary Information

### 0.1 Parameter values

#### 0.1.1 Estimating the parameters for small animal (rabbit)

1. From Google AI : 30 female offspring per year. Lifespan in the wild ie. at equilibirum in the absence of the pathogen is about 1 year. Choose max lifespan in the wild of 5 years though in captivity they can live a bit longer.
2. Rescale carrying capacity c=1.
3. Maximum lifespan 5 years gives *d*_0_ = 1/5.
4. To get natural lifespan about 1 year *d*_0_ + *k*_*d*_ ∗ *c* = 1 gives *k*_*d*_ = 1 *−* 1/5 = 0.8
5. 30 female offspring per year gives *b*_0_ = ln(30) ≈ 3.5
6. At steady-state (i.e. *c* = 1): (*b*_0_ *− k*_*b*_ ∗ *c*) = (*d*_0_ + *k*_*d*_*c*) = 1 which gives *k*_*b*_ 2.5
7. We can maintain the same eqm death rate by *d*_0_ = 0.8 *k*_*d*_ = 0.2
8. We can maintain the same eqm birth rate by *b*_0_ = 1.5 *k*_*b*_ = 0.5

#### 0.1.2 Estimating the parameters for lower homeostasis

1. We keep the same carrying capacity *c* = 1
2. Let new *b*_0_ = 1.5 and *d*_0_ = 4/5 = 0.8
3. so new *k*_*d*_ = 1 *−* 4/5 = 1/5 = 0.2
4. and new *k*_*b*_ = 1.5 *−* 1 = 0.5

Note for simulations: During the initial epidemic phase the simulations often resulted in very low number of infecteds and populations becoming negative in simulations. We thus added a very small term equivalent to constant infections from outside of the order of 10^*−*5^ as can be seen in the code.

### 0.2 Supplementary Figures

